# Comprehensive Analysis of Constraint on the Spatial Distribution of Missense Variants in Human Protein Structures

**DOI:** 10.1101/109652

**Authors:** R. Michael Sivley, Jonathan Kropski, Jonathan Sheehan, Joy Cogan, Xiaoyi Dou, Timothy S. Blackwell, John Phillips, Jens Meiler, William S. Bush, John A. Capra

## Abstract

The spatial distribution of genetic variation within proteins is shaped by evolutionary constraint and thus can provide insights into the functional importance of protein regions and the potential pathogenicity of protein alterations. Here, we comprehensively evaluate the 3D spatial patterns of constraint on human germline and somatic variation in 4,568 solved protein structures. Different classes of coding variants have significantly different spatial distributions. Neutral missense variants exhibit a range of 3D constraint patterns, with a general trend of spatial dispersion driven by constraint on core residues. In contrast, germline and somatic disease-causing variants are significantly more likely to be clustered in protein structure space. We demonstrate that this difference in the spatial distributions of disease-associated and benign germline variants provides a signature for accurately classifying variants of unknown significance (VUS) that is complementary to current approaches for VUS classification. We further illustrate the clinical utility of our approach by classifying new mutations identified from patients with familial idiopathic pneumonia (FIP) that segregate with disease.

---

Patterns of genetic variation along the human genome provide insight into functional and evolutionary constraints on different loci. A lack of common genetic variation in a locus is often indicative of functional constraint, suggesting that sequence changes negatively influence reproductive fitness^1^. The first systematic examinations of fully sequenced human genomes established consistently stronger constraint (i.e., less genetic variation) in protein-coding regions compared to non-coding sequences^2–5^; exons harbor approximately half the level of genetic variation as introns and non-coding flanking sequences. Furthermore, early candidate gene sequencing studies identified lower rates of non-synonymous variation than synonymous variation within protein-coding regions^6^, highlighting the increased constraint on protein-altering mutations. Quantifying these patterns of constraint improved the ability to identify functional regions and interpret the phenotypic effects of genetic mutations^7,8^. Building on exome-sequencing data from tens of thousands of individuals, recently-developed methods have analyzed the frequency of variation in coding regions to provide estimates of the constraint on genes based on intolerance to variation^8,9^. While these approaches have identified strongly constrained genes in which variation is likely to be pathogenic, gene-level assessment of constraint does not identify the specific protein regions and functions that are constrained and may overlook genes with such localized signatures of constraint.

Proteins are often composed of multiple domains that perform distinct functions. Constraint on missense variation varies between these regions; some regions are highly constrained while others are more tolerant of variation. Indeed, mutations to spatially distinct regions within the same protein often influence risk for different diseases^10,11^. However, spatial patterns of constraint are often visualized on an ad hoc basis by mapping sequence-derived measures of constraint into three-dimensional protein structure models^12^. Recently, structural analyses of somatic mutations from tumor samples have identified spatial clusters of mutations in several proteins^13–18^. These clusters often overlap known functional regions of oncogenes and tumor suppressors, and may assist in identifying functional driver mutations. These studies illustrate how spatial constraint on mutations can identify structural regions relevant to protein function and disease, and suggest that similar discoveries may be derived from a comprehensive analysis of germline variation. Additionally, the considerable disagreement in the clusters and proteins identified between analyses of somatic mutations suggests the need for a consistent statistical framework in which to evaluate spatial patterns of genetic variation in protein structures.

The recent abundance of human population-based sequencing studies^2,19,20^ paired with growth in the number of solved structures deposited in the protein data bank (PDB) facilitates the systematic spatial analysis of functional constraint on naturally occurring germline and somatic mutations in protein structure. In this article, we describe the comprehensive mapping of millions of human genetic variants into 4,568 solved human protein structures. We then introduce an analytical method for quantifying and comparing spatial distributions of genetic variation within protein space. This method enables us to identify significant differences in the spatial distributions of specific classes of variants— including benign and pathogenic—that reflect patterns of constraint on protein structure and function. We then describe PathProx, an algorithm for classifying the pathogenicity of missense variants of unknown significance (VUS) based on relative proximity to known pathogenic and benign variants, and demonstrate that its performance is competitive with and complementary to common pathogenicity prediction algorithms. Finally, we illustrate the clinical utility of PathProx 3D spatial analyses by classifying new mutations to the RTEL1 DNA helicase protein that segregate with disease in pedigrees with familial idiopathic pneumonia (FIP).

## Results

### Quantifying Constraint on Spatial Patterns of Genetic Variation

We mapped genetic variants from three large data sets into 4,568 human proteins with available experimentally-derived protein structures. We considered: 137,352 synonymous and 210,007 missense variants from exome sequencing of 60,706 diverse unrelated adults from the Exome Aggregation Consortium (ExAC) dataset, 4,888 missense Mendelian disease variants from the ClinVar pathogenic dataset, and 12,230 recurrent somatic missense variants (observed in at least two independent human tumor samples) from the Catalogue of Somatic Mutations in Cancer (COSMIC) dataset.

To quantify and contrast patterns of spatial constraint on different variant sets, we developed a statistic for evaluating deviations from a random spatial distribution based on

Ripley’s *K* (see Methods). Genetic variation in a protein can deviate from an unconstrained spatial distribution in two ways: it may be significantly more clustered or significantly more dispersed than expected (Figure 1a). By quantifying the density of variation around each variant in increasingly larger neighborhoods, this method identifies clustering and dispersion at any distance scale (Figure 1b). We determined the significance of observed distributions using a permutation procedure that accounts for the overall protein fold (Figure 1c-e; Methods). Finally, the method produces a Z-score-based statistic that quantifies whether the observed variants are more clustered (positive value) or more dispersed (negative value) than expected in the absence of spatial constraint (Figure 1d-e). This approach allows for direct comparison of results across structurally distinct proteins.

**Figure 1.**
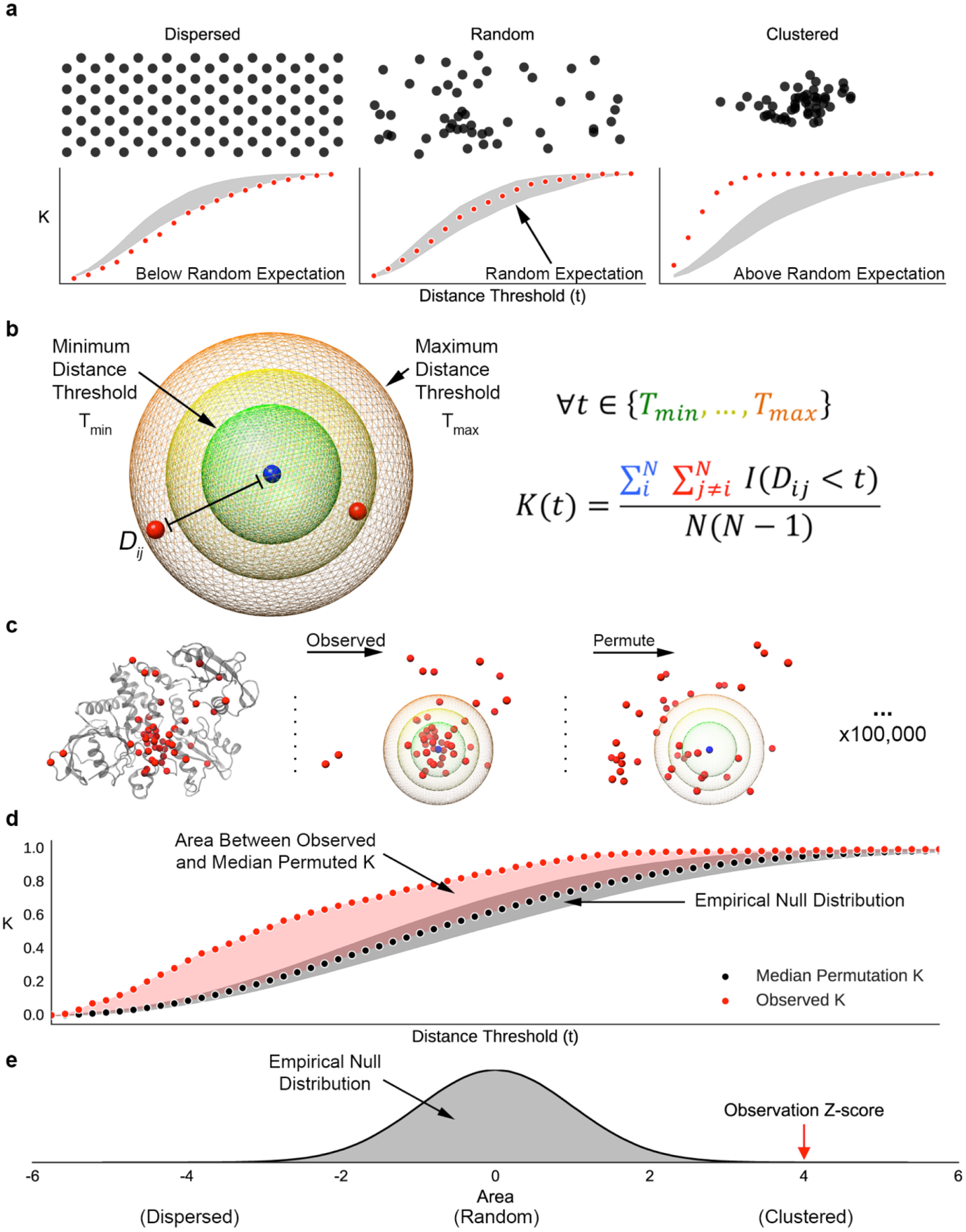
**Schematic of our framework for evaluating the spatial distribution of genetic variants**. (a) Spatial distributions can diverge from random in two ways; they may have fewer neighbors than expected by chance (dispersed) or more neighbors than expected by chance (clustered). Example distributions are illustrated in reference to a random spatial distribution in 2D. Below each set of points, the resulting K statistic at multiple distance thresholds (red) is plotted in reference to the expected K distribution under a random distribution (gray). K values below the range expected at random indicate dispersion, and K values above indicate clustering. (b) Definition of the K statistic. For a range of distance thresholds (*t*), the number of variants neighboring each variant is normalized by the total number of pairs. Two variants are considered neighbors if the distance between them (*D*_*ij*_) is less than *t. I* is an indicator function that evaluates to 1 when the condition is true and 0 otherwise. (c) The observed K values are evaluated in reference to an empirical null distribution generated from 100,000 random permutations of variant locations within the proteinstructure. (d) The spatial distribution trend for each protein is summarized by calculating the area between the observed K values (red points) and the median permuted K values (black points). (e) This process is repeated for the K values resulting from each permuted set to generate an empirical null distribution. From this distribution, we calculate a Z-score and *P*-value for the observed area. Positive Z-scores indicate clustering, negative Z-scores indicate dispersion, and Z-scores near zero indicate a lack of spatial constraint.

### Synonymous and missense variants have different spatial distributions

Synonymous genetic variants can have functional effects, e.g., by influencing gene regulation, mRNA stability, or translational efficiency; however, they rarely influence protein structural conformation^21,22^. Thus, we hypothesized that the distribution of synonymous variants in protein structure is not subject to significant spatial constraint. Consistent with this hypothesis, synonymous variants from the ExAC dataset are nearly randomly distributed in protein space (Figure 2a) and exhibit very modest clustering (median *Z* = 0.07; *P* = 0.0005, sign test). Individually, only one protein out of 4,483— Myomesin-1 (MYOM1), a long repetitive filamentous protein expressed in muscle cells—showed significant evidence of a non-random synonymous variant distribution (FDR<0.1). These results indicate that synonymous genetic variation is generally randomly distributed in the context of protein structure.

In contrast, the spatial distribution of nonsynonymous (missense) variants is likely constrained by the functional consequences of amino acid substitutions. Thus, we hypothesized that missense variants from the ExAC dataset are non-randomly distributed within protein structure. Indeed, missense variants displayed significant constraint on their spatial distribution (Figure 2b). There was a strong overall trend towards spatial dispersion (median *Z* = –0.45; *P* = 1.1x10^-106^, sign test), though we also identified instances of significant clustering in individual proteins (Figure 2b; Table S1). Comparing the observed Z-score distributions of missense and synonymous variants revealed a significant difference in their spatial patterns (Figure 2a vs. 2b; *P* = 2.71x10^-120^, Mann-Whitney *U* test).

Residues tolerant of missense variation had increased solvent accessibility compared to all residues (Figure S2, *P* ≈ 0, Mann-Whitney U test), consistent with previous analysis of missense variants from the 1000 Genomes Project^23^. Furthermore, significantly dispersed missense variants had greater solvent accessibility than missense variants overall (Figure S2, *P* = 5x10^-54^). In contrast, significantly clustered missense variants were no more or less solvent accessible than all residues. (*P* = 0.39). These results suggest that significant missense dispersion reflects constraint against substitutions in the core of a protein. Missense clustering, however, can occur at any location regardless of proximity to the protein core or surface.

**Figure 2.**
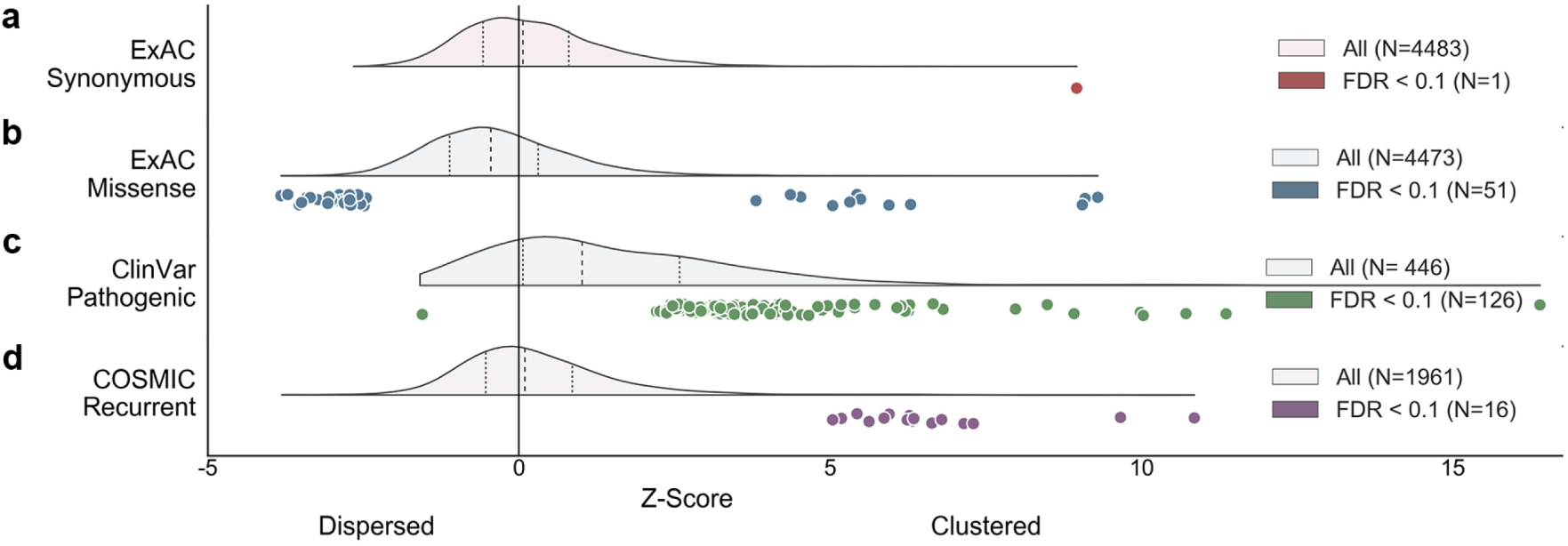
**Different types of protein-coding variants have different spatial constraints.** Each panel summarizes the spatial constraints on a different variant set. For each set, the distribution of protein-summary Z-scores over all proteins analyzed is plotted above the axis, and the Z-scores for proteins with significant constraint (FDR < 10%) are plotted with circles below the axis. Positive and negative Z-scores indicate clustering and dispersion, respectively. (a) Synonymous variants from ExAC are approximately randomly distributed, as indicated by a Z-score distribution with median near 0 (Z = 0.07). (b) In contrast, missense variants from ExAC trend towards spatial dispersion (median Z = – 0.45). (c) Pathogenic missense variants from ClinVar are the most strongly clustered variant set (Z = 1.01), with significant clustering in 126 out of 446 proteins with a sufficient number of variants. (d) COSMIC recurrent somatic missense variants show weak clustering overall (Z = 0.1), but 16proteins exhibit significant clustering. These differences in the spatial distributions of neutral and pathogenic variants suggest strong spatial constraint on protein-coding variation.

### Pathogenic missense variants are often significantly spatially clustered

Amino acids that are evolutionarily conserved across diverse species (and thus likely functional) are spatially constrained and significantly clustered within protein structure (Figure S4)^24,25^. Thus, we hypothesized that missense variants causing Mendelian disease are also likely to be spatially clustered. We evaluated this hypothesis by quantifying the spatial distribution of germline pathogenic missense variants in proteins with three or more pathogenic variants in ClinVar (Figure 2c).

ClinVar pathogenic variants are the most clustered of all variant sets analyzed (median *Z* = 1.01, *P* = 3.32x10^-31^, sign test), and 126 of 446 (28%) proteins exhibited significant clustering at FDR < 10% (Figure 2c). We also evaluated a previously curated set of missense variants from the HGMD dataset^26,27^ and observed clustering of both dominant and recessive variants (Figure S5). Dominant variants on average formed more focal clusters than recessive variants, possibly due to a greater proportion of localized, gain-of-function mutations (Supplementary Results). The frequent clustering of pathogenic missense variants underscores the spatial constraint on protein-coding variation and highlights regions in protein structures that are functionally and clinically relevant.

Several studies of tumor-derived somatic mutations have identified clusters of missense variants that are presumed to highlight structural regions important for tumorigenesis^13–18^. In contrast to germline pathogenic variants, recurrent somatic missense variants from the COSMIC database exhibit only a modest overall trend towards spatial clustering (Figure 2d; median *Z* = 0.1; *P* = 0.39, sign test). However, consistent with previous studies, a small fraction of proteins tested (16 of 1,960, 0.8%) exhibited significant clustering. This set includes many known cancer proteins^28^, including 13 proteins identified by at least one previous study of somatic mutation clustering^13,14,–18^ (Figure S6). The lack of overall clustering of recurrent somatic variants is in stark contrast to ClinVar pathogenic variant patterns and may reflect differences in spatial constraint and phenotypic effects of variation outside of the germline^10^.

### Contrasting spatial patterns of missense constraint within protein structure

Given broad evidence of spatial constraint on both putatively neutral and pathogenic variants, we contrasted the distributions of these two variant sets within individual proteins. Specifically, we evaluated whether proteins with clustering (or dispersion) of neutral variation were also likely to exhibit clustering (or dispersion) of pathogenic variation. To examine this, we plotted each protein’s Z-score for putatively neutral missense variants from ExAC against the Z-score for pathogenic missense variants from ClinVar (Figure 3a) and recurrent somatic missense variants from COSMIC (Figure 3b).

There was no significant linear relationship between ExAC-derived and ClinVar-derived Z-scores (Figure 3a; *P* = 0.71). However, their relative distributions may be informative of protein-level constraint. As expected from our comparison of the ExAC missense and ClinVar Z-score distributions (Figure 2b, c), many proteins exhibit clustering of pathogenic germline variation on a background of ExAC variant dispersion (Figure 3a, lower right). For example, pathogenic variation in Filamin-B (FLNB), aprotein that links the cellular membrane to the actin cytoskeleton, is clustered in the second calponin-homology domain, which is responsible for actin binding, while neutral variants are dispersed throughout the structure (Figure 3c). In contrast, several structures with extreme clustering of ClinVar variants also show moderate clustering of ExAC variants (upper right). We observed a notable depletion of structures with extreme dispersion from both ExAC and ClinVar analyses (lower left).

Based on these observations, we hypothesized that accounting for the background distribution of putatively neutral missense variants could identify spatial patterns unique to pathogenic variation. To test this, we calculated the *D* statistic, a bivariate form of the *K* statistic, which evaluates whether one variant set (e.g. ClinVar) is significantly more clustered (or dispersed) than another set (e.g. ExAC)^29^. Many of the outliers from the *K* analyses (especially those with clustering of ClinVar variants) were significant in the bivariate analysis. The bivariate analysis demonstrates that even when both datasets exhibit clustering, pathogenic variants are often significantly more clustered than the neutral background (Figure 3a, b). Additionally, the analysis yielded many proteins in which pathogenic and neutral distributions were significantly different, despite neither having significant univariate *K* statistics (Figure 3a). However, 21% of proteins with significant clustering of pathogenic variation were no longer significant when compared to the distribution of neutral variants. These results demonstrate that accounting for the background distribution of neutral variants enables the detection of uniquely pathogenic patterns of constraint.

There was also no significant linear relationship between the *K* statistic Z-scores for ExAC missense variants and recurrent somatic missense variants from COSMIC (Figure 3b). The most common spatial pattern was modest clustering of the COSMIC variants and slight dispersion of ExAC variants. Seven proteins harbored significant patterns of somatic constraint once accounting for neutral missense variation (Figure 3b).

Three of these have not been identified in previous analyses of cancer mutation clustering: CBL, TET2, and DICER1. CBL is an E3 ubiquitin-protein ligase, and our analyses reveal a significant cluster of recurrent COSMIC mutations in the RING-type zinc finger domain (Figure 3d). Dominant-negative germline variants in this domain impair CBL-mediated ubiquitination and decrease the ability to negatively regulate EGFR signaling^30,31^; somatic variants in this region may have similar effects.

**Figure 3.**
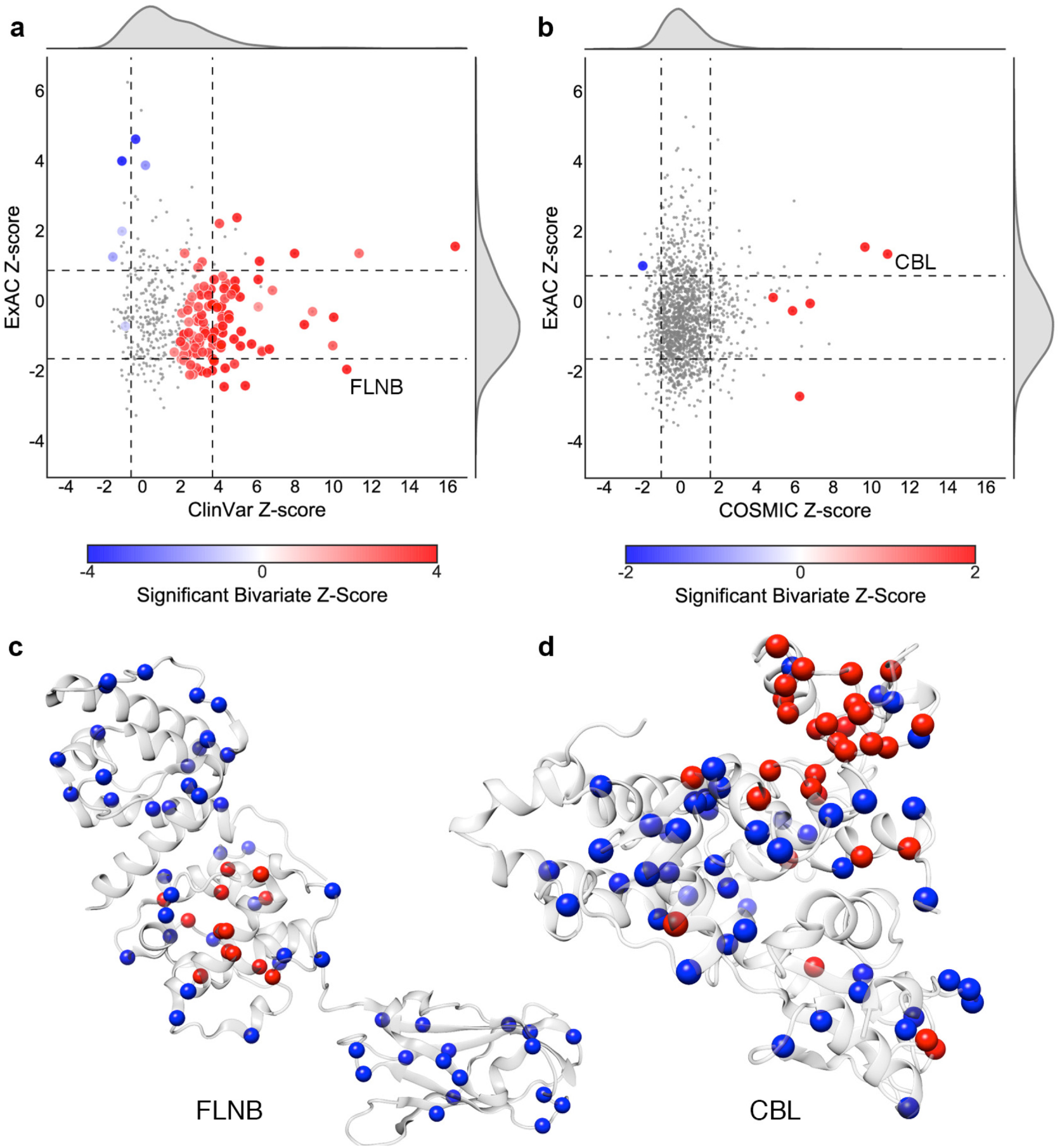
Comparison of the ExAC missense Z-scores against ClinVar pathogenic (a) and COSMIC recurrent somatic univariate Z-scores. Large circles indicate proteins with significant differences in the spatial distribution of the two sets of variants (FDR < 10%; from the bivariate analysis). Bivariate Z-scores indicate whether pathogenic/somatic variants (red) or ExAC missense variants (blue) were more clustered. The dashed lines indicate +/- one standard deviation in each dimension. (c, d) Examples of proteins with significantly different variant distributions: (c) Pathogenic variation in Filamin-B (FLNB) is clustered in the calponin-homology domain, responsible for actin binding. (d) Recurrent somatic variation in CBL, an E3 ubiquitin ligase, is clustered in the RING-type zinc finger domain, which mediates binding to E2 ubiquitin-conjugating enzymes.

### Spatial proximity to pathogenic variant clusters is predictive of pathogenicity

Given the distinct spatial patterns observed for putatively neutral and pathogenic missense variants within many proteins (Figures 2b, c and 3a), we hypothesized that accounting for the spatial distribution of known variants could improve our ability to predict the phenotypic impact of VUS. To explore this potential, we defined a simple pathogenic proximity metric (PathProx) that ranks protein residues by their relative proximity to known pathogenic and neutral missense variants (Figure 4a and Methods). We applied PathProx to 89 proteins in which ClinVar pathogenic variants were significantly clustered (*K* statistic) and significantly more clustered than ExAC missense variants (*D* statistic). In leave-one-out cross validation, PathProx performed significantly better than random (Figure 4b; median ROC AUC = 0.75, *P* = 2.4x10^-23^, sign test) and was comparable to the predictive performance of SIFT, PolyPhen2, and evolutionary conservation (*P* = 0.128, ANOVA). PathProx performance was also consistent across different protein folds as represented by CATH domains (Figure S8; *P* = 0.28, ANOVA).

Despite similar overall performance among the prediction methods (Figure 4b), on a protein-by-protein basis PathProx performance was only modestly correlated with the performance of SIFT, PolyPhen2, and evolutionary conservation (Pearson’s *r* = 0.13, 0.24, and 0.35, respectively, Figure 4c). PathProx outperformed evolutionary conservation for 35 of 89 proteins and outperformed all methods for 25 proteins (Figure 4d, Figure S9). These results demonstrate that the spatial distributions of pathogenic and neutral variants contribute distinct predictive information not captured by existing methods. However, this strong performance is contingent on clustered pathogenic variants; applying the PathProx approach without filtering for clustering yielded significantly reduced performance (N = 398, median ROC AUC = 0.55, *P* = 2.46x10^-17^, Mann-Whitney *U* test). These results suggest that proximity to pathogenic clusters, not to individual pathogenic variants, is predictive of variant pathogenicity. Thus, integrating spatial constraint with other informative features is likely to improve pathogenicity prediction.

**Figure 4.**
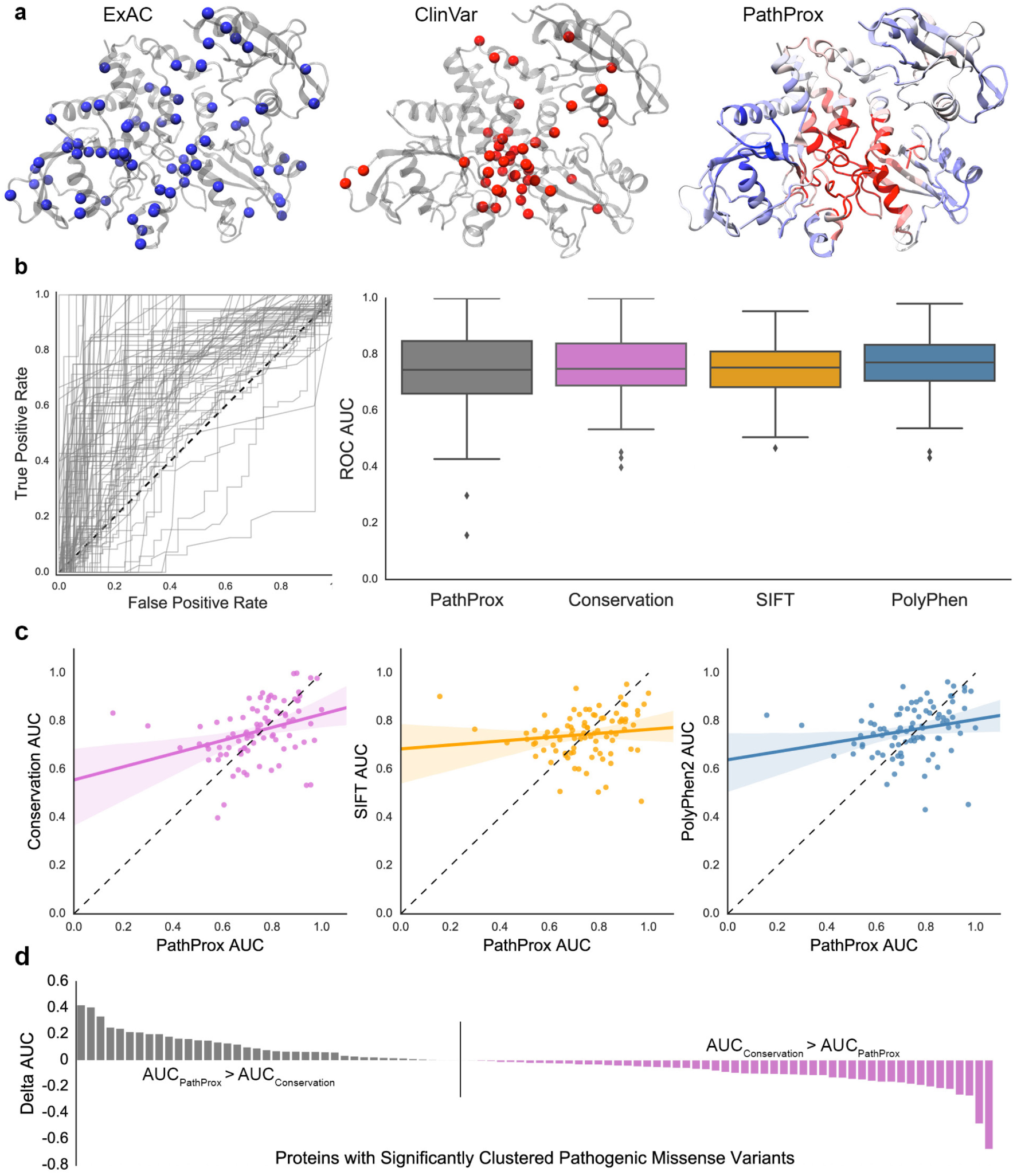
**Proximity to clusters of pathogenic variants is predictive of pathogenicity.** (a) Using ExAC (blue) and ClinVar pathogenic (red) missense variants, we calculated the pathogenic proximity (PathProx) score for all residues, indicating whether they are closer to putatively neutral (blue) or pathogenic (red) missense variants. (b) The overall predictive performance (ROC AUC) of the pathogenic proximity score was statistically comparable with other variant pathogenicity prediction methods when applied to proteins with pathogenic variant clustering. (c) Comparing the difference in performance of PathProx and other methods across proteins demonstrates that different methods perform better for different proteins. PathProx performance was only modestly correlated with evolutionary conservation (Pearson’s *r* = 0.35, *P* = 0.0008), and it was even more weakly correlated with the performance of the other methods (*r=*0.13, *P* = 0.24 for SIFT; *r* = 0.24, *P* = 0.025 for PolyPhen2). (d) PathProx outperformed (Delta AUC) evolutionary conservation on many proteins. This was true in comparisons of PathProx with all methods tested (Supplementary Figure S9) and suggests the value of integrating information about variant spatial distributions into pathogenicity prediction methods.

### PathProx Aids in the Interpretation of Missense Variants of Unknown Significance in Individuals with Familial Interstitial Pneumonia

To evaluate the practical utility of PathProx in identifying disease-causing variants in clinical applications, we applied the approach to classify missense VUS identified in patients with FIP. We first compiled 15 known pathogenic missense variants from the literature and 29 putatively neutral missense variants from the 1000 Genomes Project (Figure 5a, b). The protein structure for RTEL1 has not been experimentally determined, so we constructed two homology models covering the N and C-terminal regions of RTEL1 (Methods). Pathogenic clustering was not detected in the small C-terminal model, so we focus on the N-terminal model here and provide results for the C-terminal model in the Supplementary Results. In cross-validation using known pathogenic and putatively neutral variants, PathProx achieved a ROC AUC of 0.82 (Figure 5c). PathProx’s performance was competitive with other pathogenicity prediction methods, including SIFT and PolyPhen2.

Next, we assayed 184 FIP kindreds with targeted Sanger sequencing of *RTEL1*, a DNA helicase involved in DNA repair and telomere maintenance associated with pulmonary fibrosis^32,33^ (Supplementary Results) and identified 10 missense VUS. PathProx predicted seven of the VUS to be deleterious (Table 1). Five of these seven VUS (T55S, W512C, F559I, S688C, D719G) fully segregated with disease and were found in subjects with short telomeres in peripheral blood mononuclear cells, a biomarker of reduced *RTEL1* activity^34–36^ (Figure 5d). The other two variants (A528E, R574W) did not co-segregate with disease and were found in subjects with normal length telomeres. The three VUS predicted to be benign by PathProx (H161Q, P1107L, and F1110L) did not segregate with disease. For comparison, no other approach correctly identified all five segregating variants, and both PathProx false positive variants were also misclassified by all evaluated prediction methods (Table 1).

To explore the mechanistic basis for the association of RTEL1 mutations with disease, we mapped mutagenesis data from two studies of a homologous protein, XPD^37,38^, to our human model of RTEL1. Proximity to pathogenic variants in RTEL1 is significantly correlated with decreased ATPase activity in XPD (Figure S10; Spearman ρ=–0.62, *P* = 0.001); this suggests that pathogenic mutations in RTEL1 may similarly perturb ATPase activity in a manner that leads to disease. Detailed molecular hypotheses about how the individual segregating missense variants disrupt the structure and function of RTEL1—e.g., by disrupting protein-protein interactions (W512C) or DNA binding (F559I)—are provided in the Supplementary Results (Figure S11). These results demonstrate that the spatial proximity of missense VUS to variants of known effect can assist with VUS classification, even in the absence of a solved structure.

**Table 1.** **Pathogenicity predictions for newly identified RTEL1 missense variants of unknown significance from FIP patients.** Scores highlighted in red indicate deleterious predictions. PathProx was the only method to identify all segregating variants as pathogenic. The two non-segregating missense VUS misclassified by PathProx were also misclassified by all other prediction methods. All thresholds were applied as recommended by each method.

| Pos | Ref | Alt | Inheritance | PolyPhen2 | SIFT | ConSurf | PathProx | Model |
| --- | --- | --- | --- | --- | --- | --- | --- | --- |
| 55 | T | S | Segregating | 0.00 | 1.00 | -0.56 | 0.02 | N-terminal |
| 512 | W | C | Segregating | 0.17 | 0.48 | 0.31 | 0.35 | N-terminal |
| 559 | F | I | Segregating | 1.00 | 0.00 | -1.11 | 0.39 | N-terminal |
| 688 | S | C | Segregating | 0.91 | 0.14 | -0.62 | 0.31 | N-terminal |
| 719 | D | G | Segregating | 0.03 | 0.22 | 0.21 | 0.19 | N-terminal |
| 161 | H | Q | Non-Segregating | 0.40 | 0.16 | -0.35 | -0.08 | N-terminal |
| 528 | A | E | Non-Segregating | 0.62 | 0.05 | -0.75 | 0.23 | N-terminal |
| 574 | R | W | Non-Segregating | 0.95 | 0.00 | -0.53 | 0.08 | N-terminal |
| 1107 | P | L | Non-Segregating | 0.63 | 0.01 | NA | -0.12 | C-terminal |
| 1110 | F | L | Non-Segregating | 0.00 | 1.00 | NA | -0.15 | C-terminal |

**Figure 5.**
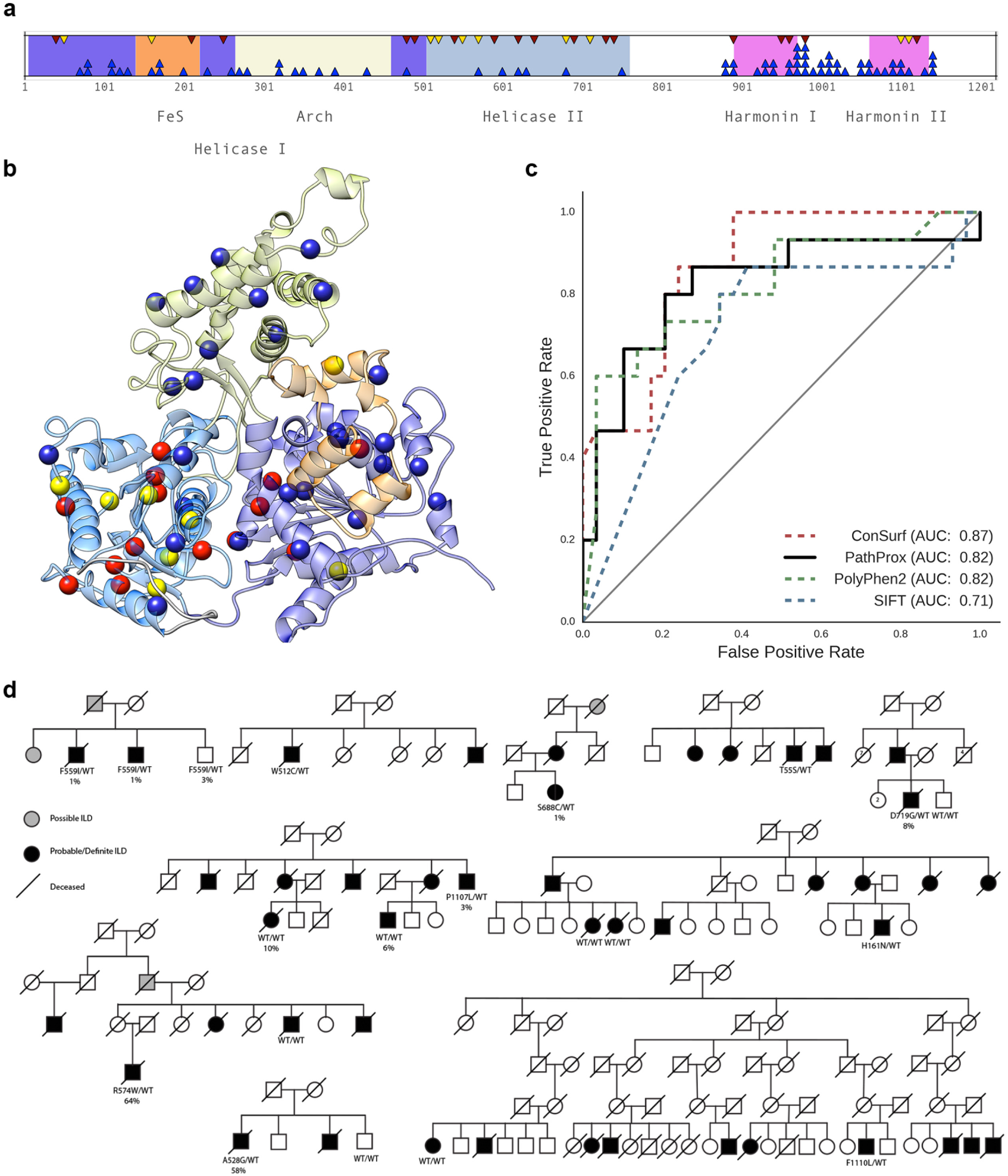
**Identification and classification of novel pathogenic familial interstitial pneumonia (FIP) variants in RTEL1.** (a) Genotyping 184 FIP patients identified 10 new missense variants of unknown significance (VUS) inRTEL1 (Table 1). The locations of ClinVar pathogenic (red), putatively neutral 1000 Genomes (blue), and new candidate FIP (yellow) missense variants are plotted in the context of the RTEL1 protein sequence and known domains. The locations of pathogenic, putatively neutral, and candidate variants in the RTEL1 N-terminal structural model. Leave-one-out cross validation of PathProx applied to characterized RTEL1 variants yielded a comparable area under the ROC curve (AUC) to PolyPhen2, improved on SIFT, and was outperformed by evolutionary conservation scores. (d) Analysis of pedigrees of FIP patients demonstrated that five VUS segregate with disease. PathProx provided the most accurate classification of these RTEL1 VUS (Table 1). These results demonstrate how considering the spatial distribution of known pathogenic and neutral variants can identify pathogenic hotspots and assist in the classification of VUS.

## Discussion

By projecting missense variation into the three-dimensional context of experimentally-derived protein structures, we quantified patterns of spatial constraint on genetic variation within human proteins. As expected, synonymous variants are nearly randomly distributed within protein structures. Reflecting the diversity of constraints on protein structure and function, missense variants exhibit significant dispersion in some proteins and significant clustering in others. However, variant-intolerance in core buried regions leads to a general trend towards spatial dispersion. In contrast, more than three quarters of proteins with ClinVar pathogenic variation display some degree of spatial clustering, and over a quarter exhibit significantly more clustering than expected in the absence of constraint. These clusters highlight protein regions that are particularly sensitive to amino acid substitutions, and capture spatial relationships that may be missed by traditional methods.

Motivated by previous analyses of tumor-derived somatic variation^13–18^, we also evaluated spatial patterns of constraint on COSMIC recurrent missense variants. As expected, we found several proteins with significant clustering of COSMIC variation, including known cancer proteins. However, in general, the somatic variants were under little spatial constraint; only 15 of 1,960 proteins tested had significant deviations from a random distribution. Several factors likely contribute to the weaker clustering of the cancer mutations compared to germline pathogenic mutations. It may reflect fundamental differences in variant tolerance between germline and somatic contexts. Germline variants are present in all tissues and are constrained to be compatible with proper development. In contrast, somatic variants influence only a subset of tissues, and thus may be tolerated in contexts that would be lethal in the germline^10^. However, technical factors may also contribute to this difference. Even though we limited our analyses to recurrent COSMIC mutations seen in multiple tumors, the set is likely is a mixture of driver and passenger mutations, and thus maybe less enriched for pathogenic mutations than the germline set.

Our direct comparison of the spatial distributions of pathogenic and neutral variation within individual protein structures identified many regions tolerant and intolerant of missense variation. From this analysis, we discovered that proteins fall on a continuum of relative constraint; for example, FLNB has a single cluster of pathogenic variants on a background of widely dispersed neutral variants, while CBL exhibits clustering of both pathogenic and neutral variants within different regions of the protein. This result has important implications for the estimation of constraint. First, across all proteins, the spatial distribution of neutral missense variants is often different than the distribution of pathogenic variation. Second, directly comparing the spatial distributions of neutral and pathogenic variation reveals patterns that are not evident from examining either pathogenic or neutral variation alone, including uniquely pathogenic patterns of spatial constraint. For example, in 21% of protein structures with significant clustering of pathogenic variants, the pathogenic variants were not significantly more clustered than neutral missense variants in the same protein. The remaining 79% of proteins exhibit clustering of pathogenic variation that cannot be explained by the general patterns of missense variation within the structure.

Knowledge of constrained regions within proteins structures is informative for the identification of pathogenic variants. Indeed, the differences we discovered in the spatial distributions of disease-causing and non-pathogenic variants can be exploited to predict variant pathogenicity. Our PathProx metric performs competitively with other common pathogenicity prediction methods on proteins with a sufficient number of characterized variants, yet its predictions are only modestly correlated with existing methods. This suggests that the spatial relationships between variants provide unique insights into variant pathogenicity. The clinical utility of this approach is illustrated by the case study in which we accurately classify RTEL1 VUS that co-segregate with disease in FIP-affected pedigrees.

The spatial patterns we detect are influenced by differences in evolutionary constraint and provide a unique perspective from which to study protein function and the phenotypic effects of coding variation. However, there are limitations to our approach. First, the utility of PathProx requires protein structural information. Solved structures are available for approximately 25% of human proteins; however, the success of PathProx on RTEL1 highlights the potential of using homology-based computational models for genes without experimentally-derived protein structures. Homology-based models are available for more than 80% of human protein structures. Second, our analyses cannot be applied to proteins that lack known variants with characterized effects. We anticipate that mapping mutations across homologous protein families, and potentially even from model organisms, will significantly increase the number of human proteins with sufficient mutations with known effects. Nonetheless, while spatial information may not be available in all contexts, the information captured is largely independent from the pathogenic signatures used by other methods, some of which incorporate structural features of single variants. Thus, there is great potential for the incorporation of the spatial distribution of variants into existing pathogenicity prediction algorithms.

In summary, our work provides a consistent statistical framework in which to identify significantly constrained spatial regions in protein structures and demonstrates significant differences in the spatial distribution of synonymous, non-synonymous, and disease-associated protein-coding variation. To facilitate further analyses, we provide ASTRID (http://astrid.icompbio.net/), a web-interface for viewing the structural locations of all ExAC, ClinVar, and COSMIC variants, along with the results of all spatial and predictive analyses, in a representative set of 4,568 solved human protein structures.

## Online Methods

### Genetic Variant and Protein Structure Datasets

We analyzed single-nucleotide variants (SNV) from Exome Aggregation Consortium^20^ (ExAC) r0.3, ClinVar (01-07-2016), and COSMIC version 74. Variant consequences and annotations were determined using v82 of the Ensembl Variant Effect Predictor for genomic build GRCh37^39^. Missense variants from the 1000 Genomes Project^19^ were use to supplement known benign variants for evaluation of variants of unknown significance in RTEL1. Synonymous SNVs in ExAC were included for comparison with ExAC missense SNVs. All other datasets were filtered to include only missense SNVs.

Genetic variants where mapped into representative protein structures through Ensembl^40^ transcripts, which were matched with UniProt^41^ accession and Protein Data Bank^42^ (PDB, 01-07-2016) IDs using cross-reference tables provided by UniProt. Reference protein sequences were aligned with observed sequences in the PDB using SIFTS^43^. Discrepancies were corrected by Needleman-Wunsch pairwise alignment with Biopython^44,45^. Proteins were represented by the subset of minimally overlapping PDB structures described by Kamburov *et al.*
^14^.

### Quantifying and comparing the spatial distributions of protein-coding mutations

We developed a framework for evaluating hypotheses about the spatial distributions of genetic variants in protein structures based on Ripley’s *K*, a spatial descriptive statistic commonly used in ecology and epidemiology^29,46,47^. The univariate (single dataset) *K* quantifies the spatial heterogeneity of a set of variants by comparing the proportion of variants within a given distance from one another to the expectation under a random spatial distribution. Variants are considered clustered if the proportion of neighbors exceeds expectation and dispersed if the number of neighbors is lower than the expectation. *K* is calculated across a range of distance thresholds (*t*), enabling the identification of clustering or dispersion at any scale (Figure 1a). We define *K* as

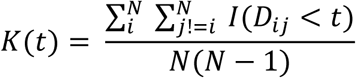

Where *N* is the number of variants in the protein structure, *Dij* is the Euclidean distance between variants *i* and *j*, and *I* is an indicator function that evaluates to 1 when *Dij* is less than the distance threshold *t* and 0 otherwise. *N(N-1)* is a normalization factor for the number of variant pairs; as a result, *K* can be interpreted as the proportion of variant pairs within distance *t* of one another. This normalization also allows for comparison between proteins with different variant counts. Variant positions are defined as the centroid of the reference amino acid (Figure 1b).

Missense variants are constrained to the positions of amino acids in a protein structure, so complete spatial randomness is not a valid null model for randomly distributed variants (Figure 1c). To account for these constraints, we calculate the empirical null distribution of *K* through 100,000 random permutations of variant positions within the structure. Two-tailed *P*-values are derived from the proportion of permuted *K* values more extreme than the observed *K* value. Z-scores are calculated to quantify the direction (clustering or dispersion) and magnitude of the effect.

To evaluate the spatial distribution of real-valued attributes (e.g., evolutionary conservation, solvent accessibility) we compute a weighted form of the univariate analysis. We define the weighted *K* as

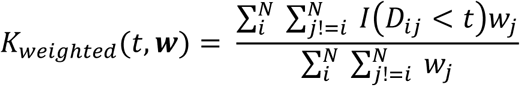

Where *wj* is the weight associated with protein position *j*. We use this approach to assesses whether the weights exhibit significant spatial constraint (clustered or dispersed) beyond what is explained by position. Thus, we evaluate the significance of the weighted *K* by permuting the weights over the fixed amino acid positions and empirically computing *P*-values. When *N* is small, the ability to assess the significance of the weighted *K* is limited by the number of unique permutations.

This framework can quantify spatial distributions across a range of distances, capturing clustering or dispersion at any scale. To focus on spatial patterns at biologically relevant distance scales, we evaluated distances between the minimum observed distance between variants to half the maximum observed distance between variants. In proteins for which the minimum distance is greater than half the maximum distance, we extend the range to the maximum distance. The distance threshold that yields the most extreme Z-score marks the scale at which the spatial distribution differs most from expectation.

To summarize spatial patterns into a protein-level summary statistic, we computed the area between the observed K curve and the median empirical null K curve using Simpson’s rule (Figure 1d). This summarization captures the general spatial tendencies for each protein. This process is repeated for each permuted K-curve to generate an empirical null distribution. From this distribution, we calculate a permutation *P*-value and Z-score for the observed area (Figure 1e). Positive Z-scores indicate clustering, negative Z-scores indicate dispersion, and Z-scores near zero indicate spatial randomness (e.g. a lack of spatial constraint). We controlled the False Discovery Rate (FDR) at 10% by computing q-values from the protein-summary *P*-value distribution in each analysis^48^ (github.com/nfusi/qvalue).

### Bivariate D for spatial comparisons between variant datasets

The univariate *K* quantifies constraint on the spatial distribution of a single set of variants, but many biological questions involve comparisons between variants of different types (e.g., neutral and pathogenic). We adapted the bivariate *D* statistic^29^ to enable these comparisons. The bivariate analysis evaluates whether one set of variants is more or less clustered than another by computing the difference in their univariate *K*.

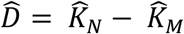

Where N and M are two sets of variants. Similar to the weighted univariate *K*, the bivariate *D* evaluates whether the process by which dataset labels (e.g. pathogenic, neutral) are determined is spatially constrained. We determined the Z-score and significance of the bivariate *D* through random permutation of the class labels over fixed variant positions.

### Relative proximity to pathogenic variation as a predictor of pathogenicity

To measure the average proximity of a variant *x* to a set of known variants *Y*, we identify the proportion of variants in *Y* within some distance of *x*. The average proximity of *x* to *Y* is defined as

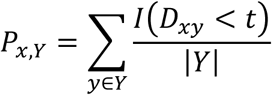

Where *I* is an indicator variable that evaluates to 1 if variants *x* and *y* are within distance *t* and *|Y|* is the number of variants in dataset *Y*. For each protein structure, we selected the distance threshold at which the bivariate Z-score between pathogenic and neutral variants was most extreme, indicating the distance at which their spatial distributions are most different. The pathogenic proximity (PathProx) score for each variant was then defined as the difference in average proximity to pathogenic and neutral variation.

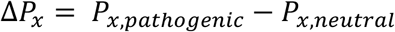

Such that values of ∆ greater than 0 indicate that variant *x* has greater spatial proximity to pathogenic variants than neutral variants.

To evaluate the ability of the pathogenic proximity score to classify pathogenic and neutral mutations, we performed leave-one-out cross validation of ClinVar pathogenic and ExAC missense variants in proteins for which ClinVar pathogenic variants were determined to be significantly clustered by both univariate and bivariate analyses. We then calculated receiver-operator-characteristic (ROC) and precision-recall (PR) curves from the pathogenic proximity score of each variant and summarized predictive performance by computing the area under the ROC and PR curves (AUC). We used Analysis of Variance (ANOVA) to compare pathogenic proximity performance with evolutionary conservation^49^ and pathogenicity scores from SIFT^50^ and PolyPhen2^51^. Default thresholds for each method were used in predicting variant pathogenicity.

### Subjects and Specimens

The Institutional Review Boards from Vanderbilt University, Duke University, University of Colorado and National Jewish Hospital approved this investigation and subjects or their surrogates provided written informed consent prior to enrollment in the study. Subjects were identified from the Familial Interstitial Pneumonia (FIP)/Familial Pulmonary Fibrosis (FPF) registries at Vanderbilt University, the University of Colorado, and National Jewish Hospital^32^. FIP was defined by the presence of Idiopathic Interstitial Pneumonia (IIP) in two or more family members, including Idiopathic Pulmonary Fibrosis (IPF) in at least 1 individual. Phenotypes of subjects selected for sequencing were ascertained using ATS/ERS criteria for IIP^52^. The affected status of deceased individuals was determined by review of available medical records, autopsy material, or by death certificates. DNA was isolated from blood and/or paraffin-embedded lung tissue using a PureGene Kit (Gentra Systems, Minneapolis, MN). Whole exome sequencing and Sanger sequencing of *RTEL1* in extended pedigrees and probands from FIP kindreds was performed as previously described^32^.

### Constructing the Structural Model of RTEL1

We applied nine computational modeling algorithms to the RTEL1 protein sequence: GeneSilico^53^, HHpred^54^, I-TASSER^55^, M4T^56^, Pcons5^57^, Phyre2^58^, RaptorX^59^, Robetta^60^, and SWISS-MODEL^61^. RaptorX produced the highest-coverage model, which consisted of two well-folded domains spanning residues 1–769 and 881–1151. This model was based on seven PDB structures: 4a15^62^, 3crv^38^, 2fi7^63^, 2gm7^64^, 4pjq^65^, 2vrw^66^, 4a64^67^. To improve quality, the model was relaxed using Rosetta version 2015.19^68^, then subjected to 1000 rounds of loop modeling^69^ using perturb_kic_with_fragments. This new structural model of RTEL1 is available as Supplementary File 1.

